# Nitrogen signaling factor triggers a respiration-like gene expression program

**DOI:** 10.1101/2023.12.18.572203

**Authors:** Shin Ohsawa, Michaela Schwaiger, Vytautas Iesmantavicius, Rio Hashimoto, Hiromitsu Moriyama, Hiroaki Matoba, Go Hirai, Mikiko Sodeoka, Atsushi Hashimoto, Akihisa Matsuyama, Minoru Yoshida, Yoko Yashiroda, Marc Bühler

## Abstract

Microbes have evolved intricate communication systems that enable individual cells of a population to send and receive signals in response to changes in their immediate environment. In the fission yeast *Schizosaccharomyces pombe*, the oxylipin Nitrogen Signaling Factor (NSF) is part of such communication system, which functions to regulate the usage of different nitrogen sources. Yet, the pathways and mechanisms by which NSF acts are poorly understood. Here, we show that NSF physically interacts with the mitochondrial sulfide:quinone oxidoreductase Hmt2 and that it prompts a change from a fermentation- to a respiration-like gene expression program independently of the carbon source. Our results suggest that NSF activity is not restricted to nitrogen metabolism alone and that it could function as a rheostat to prepare a population of *S. pombe* cells for an imminent shortage of their preferred nutrients.

## INTRODUCTION

Unicellular organisms must be able to sense and react to changes in their immediate environments to maximize growth and survival. Classic examples are acute responses to starvation or other stressful conditions such as osmotic, oxidative, or temperature stress, heavy metals, or DNA damage. Besides shortage of nutrients, they must also accommodate changes in the abundance of nutrients that they favor. For this, catabolite repression (CR) strategies are utilized by various species of bacteria and fungi, which enables them to preferentially utilize high-quality nutrients (Nair & Sarma, 2021). By modulation of CR, they can accommodate nutritional changes in their environment.

Carbon and nitrogen are the main energy sources for sustaining biosynthetic processes and must be taken up in large quantities from the environment. Different species have preferences for certain carbon and nitrogen sources that they rapidly metabolize to generate energy for growth and niche colonization. In the presence of these favored energy sources, utilization of less preferred carbon and nitrogen sources is repressed, phenomena referred to as carbon and nitrogen catabolite repression (CCR and NCR, respectively). Because of the impact CCR and NCR have on virulence of human pathogens, a good mechanistic understanding of carbon and nitrogen metabolism in microbes is of great biomedical importance (Ries *et al*, 2018). It is also highly relevant for industrial applications that rely on microorganisms for the generation of valuable bio-products (Nair & Sarma, 2021).

The selective usage of one energy source over another is generally considered to be transient, acute, and cell autonomous. However, recent studies revealed the existence of chemical communication between cells that transforms metabolic responses (Jarosz *et al*, 2014; Takahashi *et al*, 2012). This can occur between cells of the same or across species, or even kingdoms, as exemplified by the bacterial metabolite lactic acid, which serves as a diffusible signal that enables neighboring *Saccharomyces cerevisiae* cells to bypass CCR (Jarosz *et al*, 2014; Garcia *et al*, 2016). An intra-species chemical communication system that regulates NCR was discovered in the fission yeast *Schizosaccharomyces pombe* (Sun *et al*, 2016; Takahashi *et al*, 2012). Here, uptake of the branched-chain amino acids (BCAA) isoleucine (Ile), leucine (Leu), and valine (Val) is suppressed in the presence of high-quality nitrogen sources such as ammonium or glutamate, because the expression of transporters or permeases that are needed for the uptake of poorer nitrogen sources are down- regulated (Zhang *et al*, 2018). Thus, *S. pombe* cells rely on their own BCAA synthesis to sustain growth. Therefore, auxotrophic mutants, such as Leu-auxotrophic *leu1-32* or BCAA- auxotrophic *eca39Δ* cells, are unable to grow on minimal medium containing a high-quality nitrogen source, even when supplemented with BCAA (Takahashi *et al*, 2012; Sun *et al*, 2016). Notably, though, growth of *leu1-32* and *eca39Δ* cells can be restored when prototrophic cells are plated adjacent to the mutant cells. This adaptive growth is not observed near *S. cerevisiae* cells (Takahashi *et al*, 2012; Sun *et al*, 2016). This intriguing observation suggested that the prototrophic cells secrete a diffusible molecule that revokes NCR in the auxotrophic cells. Indeed, the fatty acids 10(*R*)-acetoxy-8(*Z*)-octadecenoic acid and 10(*R*)-hydroxy-8(*Z*)-octadecenoic acid were found to be secreted by the prototrophic cells, now referred to as Nitrogen Signaling Factors (NSFs). Importantly, synthetic NSFs are sufficient to bypass NCR, i.e. *leu1-32* and *eca39Δ* cells can grow on minimal medium containing a high-quality nitrogen source when supplemented with BCAA and synthetic NSF (Sun *et al*, 2016). Thus, NSFs are part of a species-specific communication system that enables *S. pombe* cells adapting to changing nutritional conditions by evading NCR. However, the pathways in which NSFs function in such adaptation are not known.

Here, we employed forward genetics, genomics, and chemical biology approaches to gain insights into NSF-mediated adaptive growth of *S. pombe* cells. We show that mitochondrial respiration counteracts NCR, that NSF exposure triggers a change from a fermentation- to a respiration-like gene expression program, and that the mitochondrial sulfide:quinone oxidoreductase Hmt2 is a direct target of NSF. Thus, NSF is a transmissible signal that turns on mitochondrial respiration to maximize growth in response to changes in the availability of nutrients.

## RESULTS

### Identification of genes that are required for NSF-mediated adaptive growth

Because cellular uptake of BCAA is suppressed in the presence of high ammonium concentrations, *leu1-32* auxotrophs are unable to grow unless they are exposed to a diffusible signal such as NSF that is secreted by neighboring prototrophs (Figure 1A). To identify genes that are required for NSF-mediated adaptive growth, we decided to perform genetic screens using the BIONEER haploid *S. pombe* Genome-wide Deletion Mutant Library that covers 3420 non-essential gene knock-out strains (Kim *et al*, 2010) (Table. S1). Notably, this library harbors auxotrophic mutations not only in *leu1^+^*, but also in the *ura4^+^* and *ade6^+^* genes (*leu1-32, ura4-D18*, and *ade6-M210* or *M216*). To exclude potential confounding effects that might stem from the *ura4^+^* and *ade6^+^* auxotrophic alleles (Takahashi *et al*, 2012), we first crossed the *h*^+^ BIONEER library with a *h^-^* strain that had an auxotrophic mutation in the *leu1^+^* gene only and a nourseothricin resistance marker downstream of the *mat1-Mc* gene (*leu1-32 mat1<<natMX,* Figure 1B). From these crosses, we could recover 3225 individual *h^-^*knock-out strains that are auxotrophic for leucine specifically (Leu-auxotrophic knock-out library; Figure 1B, Table S1).

**Figure 1.**
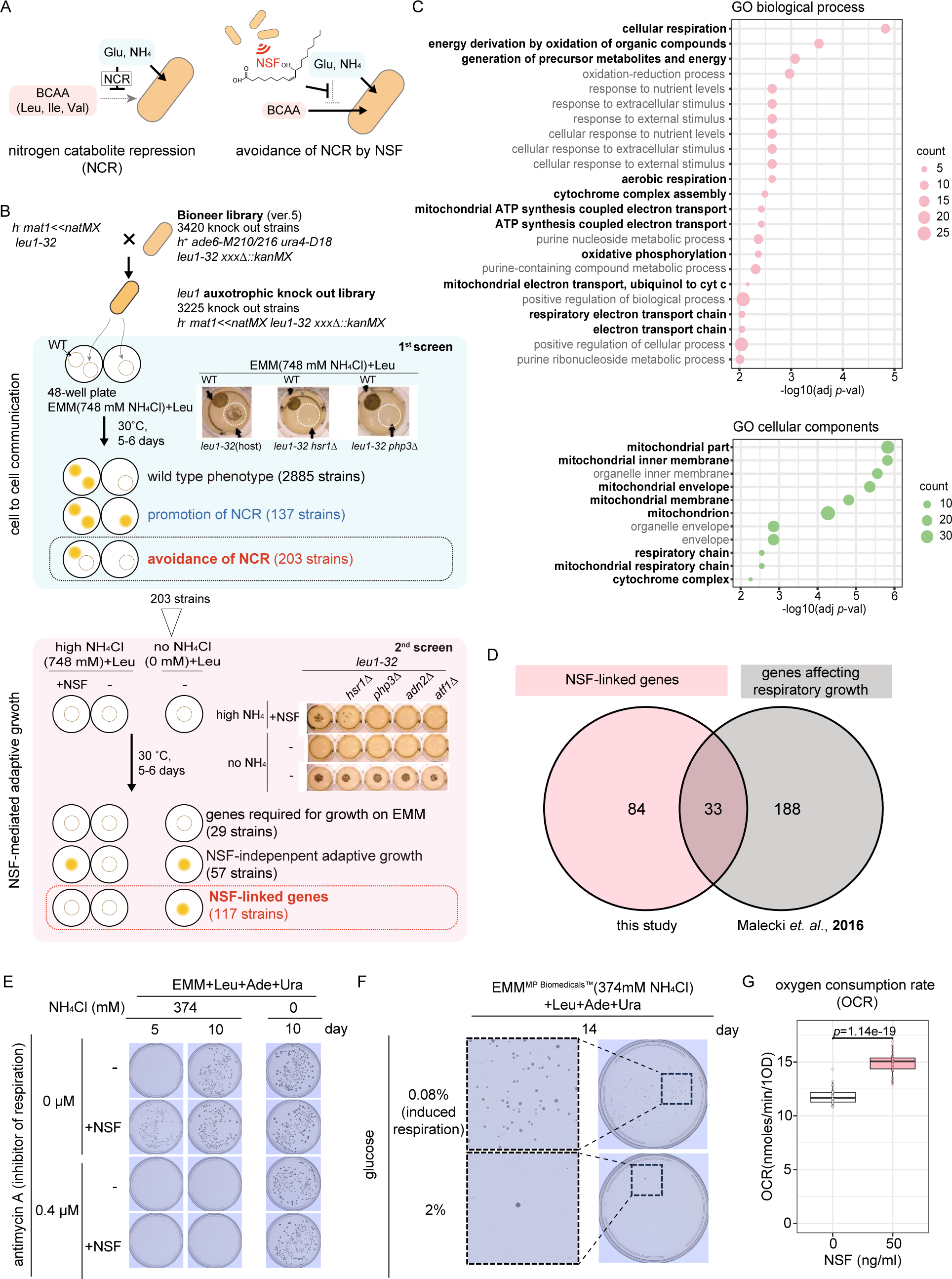
NSF-mediated adaptive growth is linked to respiration. **(A)** Scheme illustrating NCR. Cellular uptake of branched-chain amino acids (BCAA) is suppressed in the presence of glutamate (Glu) or ammonium (NH_4_) (left). This is bypassed upon exposure to NSF (right). **(B)** Outline of the genetic screens performed in this study. **(C)** GO enrichment analyses of NSF-linked genes by AnGeLi (Bitton *et al*, 2015). GO terms linked to mitochondria are written in bold letters. **(D)** Venn diagram showing the overlap of NSF- linked genes revealed by this study and genes that were previously shown affect respiratory growth (Malecki & Bähler, 2016). **(E)** NSF-mediated adaptive growth under high ammonium conditions when respiration is inhibited. **(F)** Cellular growth under high ammonium conditions when respiration is induced by low glucose concentrations (0.08%). **(G)** Oxygen consumption rate of cells that were exposed to synthetic NSF. *p*-value was calculated using a two-sided t-test.

We performed a first screen to identify those genes that are required when auxotrophs must communicate with prototrophs to sustain growth under high ammonium conditions (cell-to-cell communication, CCC). For this, the 3225 Leu-auxotrophic knock-out strains were spotted in 48-well high ammonium agar plates (EMM(748mM NH_4_Cl)+Leu) either alone, or in the vicinity of a wild type (*wt*) prototroph. This revealed three different growth phenotypes (Figure 1B, middle): i) 2885 mutants that grew only in the presence of *wt* prototrophs (wild type phenotype; Table. S1), ii) 137 mutants that grew irrespective of neighboring *wt* prototrophs (genes required for NCR; Table S1), and iii) 203 mutants that failed to grow either alone or in the presence of neighboring *wt* prototrophs (genes required for the avoidance of NCR; Table S1). Thus, we have identified 203 genes that are necessary for *leu1- 32* auxotrophs to respond to signals, sent out by *wt* prototrophs, that enable them to take up leucine from the environment (avoidance of NCR).

Although NSF has been identified as a diffusible signaling molecule that is sufficient to sustain the growth of leucine auxotrophs under high ammonium conditions (Sun *et al*, 2016), it remains unknown whether other signals are secreted that could function redundantly to NSF. To test how many of the 203 genes required for the avoidance of NCR would be specifically required for NSF-mediated adaptive growth, we performed a second screen (Figure 1B, bottom). For this, the 203 mutant strains were spotted on EMM(748mM NH_4_Cl)+Leu plates coated either with synthetic NSF (10(*R*)-hydroxy-8(*Z*)-octadecenoic acid) (Sun *et al*, 2016) or methanol as a mock control. The strains were also spotted onto plates lacking ammonium (EMM(0 mM NH_4_Cl)+Leu) to check abilities of growth on EMM, and leucine uptake and utilization. As expected, none of the strains grew on high ammonium plates (EMM(748mM NH_4_Cl)+Leu). Of those, 29 mutants grew under neither condition tested, indicating that these genes are related to EMM growth, leucine uptake, or utilization of ammonium as the sole nitrogen source (“genes required for growth on EMM” in Figure 1B). Growth on high ammonium plates was rescued for 57 mutants in the presence of NSF (NSF-independent adaptive growth in Figure 1B). That is, these genes are not needed to respond to NSF. Yet, because they are needed for adaptive growth (1^st^ screen), they are likely required to respond to a transmissible signal that is different from NSF (Sun *et al*, 2016; Chiu *et al*, 2022). This was not the case for 117 mutants, which grew normally without ammonium, but not at high ammonium concentrations despite the addition of NSF. Thus, we have identified 117 genes that are required for NSF-mediated adaptive growth (referred to as “NSF-linked genes” listed in Tables S1 and S3).

### Mitochondrial respiration counteracts NCR

Analyzing the list of 117 NSF-linked genes with AnGeLi (Bitton *et al*, 2015), we noticed enrichment of genes that are encoding proteins that are related to mitochondrial respiration (Figure 1C and Table S2). Interestingly, 33 NSF-linked genes have been previously implicated in respiratory growth when glycerol serves as carbon source (Malecki *et al*, 2016) (Figure 1D and Table S3). This suggests that NSF-mediated adaptive growth depends on mitochondrial respiration. To test this directly, we assessed NSF-mediated adaptive growth under high ammonium conditions by the addition of antimycin A, which is a potent mitochondrial electron transport inhibitor and can thus be used to block respiration (Heslot *et al*, 1970; Malecki *et al*, 2016). As shown in Figure 1E, antimycin A blocked NSF-mediated adaptive growth in the presence of high ammonium concentrations. In the absence of ammonium, 0.4 µM antimycin A had no effect on cell growth. *Vice versa,* when we induced respiration by lowering the glucose concentration (0.08%) (Takeda *et al*, 2015), NSF even became dispensable for growth under high ammonium conditions (Figure 1F). To check whether NSF treatment stimulates respiration, we measured oxygen consumption rates (OCR). Indeed, we observed an increased OCR after 8 hours NSF treatment (Figure 1G). From these results we conclude that mitochondrial respiration counteracts NCR, and that this is positively influenced by NSF signaling.

### Changes in gene expression upon exposure to NSF

To investigate global changes in gene expression in response to NSF exposure, we performed total RNA sequencing with cells that were grown in the presence of high ammonium concentrations and exposed to synthetic NSF for two, four, or six hours. Already after 2 hours exposure to NSF, we observed 98 and 74 genes that were significantly up- or downregulated, respectively (Figure 2A). This remained largely unchanged when cells were exposed to NSF for longer (Figure S1). Thus, NSF-induced changes in gene expression can be observed within the first 2 hours already. Of the 117 NSF-linked genes described above, only *hsr1^+^* was differentially expressed upon NSF exposure. Expression of respiration-related genes that were revealed by the screen remained unchanged. Yet, we noted that global differential gene expression patterns between NSF-treated cells (this study) and cells that had been shifted to respiration by glycerol feeding (Malecki *et al*, 2016) correlated positively (Figure 2B). Interestingly, genes previously shown to be upregulated in respiratory conditions tended to be upregulated in response to NSF exposure as well (Figure 2C). Likewise, genes downregulated in respiratory conditions appeared to be less expressed also in cells exposed to NSF (Figure 2C). These results reveal that NSF triggers a change from a fermentation- to a respiration-like gene expression program. GO enrichment analysis (Table S2) of the 92 downregulated genes revealed an overrepresentation of GO terms related to flocculation or adhesion (Figure 2D). The 156 upregulated genes were enriched for GO terms related to cellular fusion or mating, trehalose synthesis, or detoxification of ROS. For example, Catalase (Ctt1) and glutathione peroxidase (Gpx1) function as ROS scavengers in mitochondria (Paulo *et al*, 2014). Coherent with the induction of a respiration-like gene expression program, the trehalose synthesis pathway is also upregulated under respiratory conditions, in which trehalose functions as an antioxidant (Malecki *et al*, 2016). Enrichment of these GO terms thus suggests that NSF contributes to the induction of expression of antioxidants for ROS that is generated by the respiration pathway in mitochondria. We find it interesting that two major cell-cell adhesion events, mating and flocculation, are oppositely enriched in up- and down-regulated genes, respectively. Because flocculation enhances mating efficiency (Goossens *et al*, 2015), we envision that the downregulation of flocculation/adhesion-related genes by NSF could serve to avoid undesirable mating during growth.

**Figure 2.**
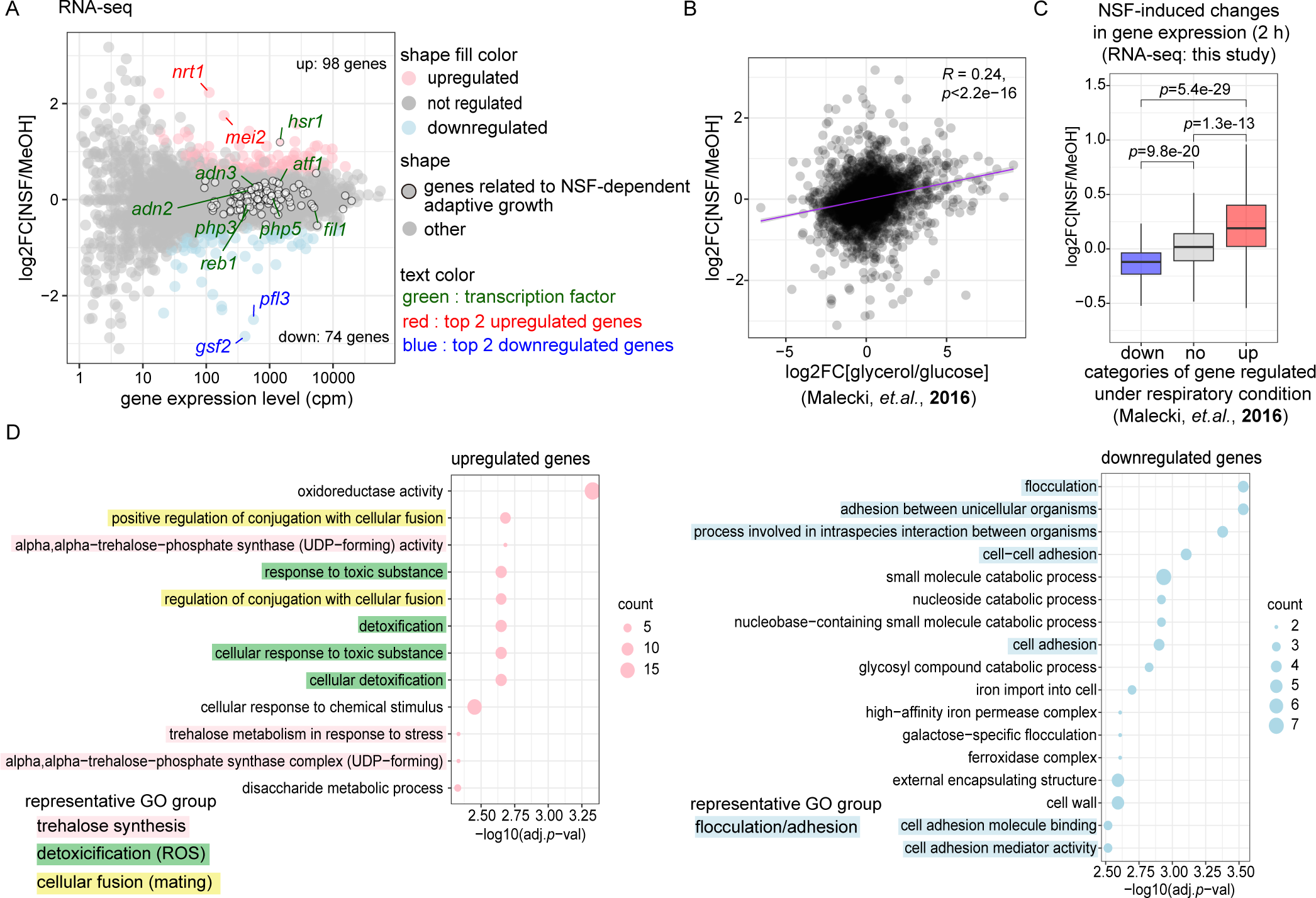
NSF-induced changes in gene expression. **(A)** MA plot showing differential gene expression (log2FC) in cells treated with MeOH or NSF for 2 hours (y-axis). x-axes denotes total transcript abundance in counts per million (cpm) in both conditions. *p*-values were calculated using the wald test and adjusted with the Benjamini and Hochberg (BH) method. Up- or down-regulated genes (FC >1.5 and adjusted *p*-value <0.05) are highlighted in pink or blue, respectively. NSF-linked genes revealed by the genetic screen are marked with a black outline. The names of the top two upregulated and downregulated genes are written red and blue, respectively. Green color was used to label the genes encoding transcription factors. **(B)** Scatter plot comparing log2FC gene expression changes induced by NSF (y-axes, this study) or glycerol feeding (x-axes)(Malecki *et al*, 2016). **(C)** Box plot showing logFC distribution of NSF-induced gene expression changes, grouped by gene expression changes induced by respiration (Malecki *et al*, 2016). *p*-values were calculated using a two-sided t-test and were adjusted using the holm method. **(D)** GO enrichment analysis of genes that are differentially expressed upon NSF exposure. GO term enrichment is shown on the x-axis (as -log10 adjusted *p*-value) for all GO terms enriched (adj.*p*-val < 0.05) among upregulated (left) or downregulated (right) genes. Representative GO groups are highlighted with indicated color.

### The NSF-responsive amino acid transporter 1 gene

Of the genes with a gene expression level higher than 100 cpm, *SPBPB2B2.01^+^*and *mei2^+^* were the two most highly upregulated genes, whereas *gsf2^+^* and *pfl3^+^* were the two most strongly downregulated genes (Figure 2A). Further analyses revealed that expression of all four genes changed in an NSF dose-dependent manner (Figure 3A). Notably, *SPBPB2B2.01^+^* is an uncharacterized gene whose protein product has an inferred biological role in amino acid transmembrane transport (https://www.pombase.org/gene/SPBPB2B2.01). This is interesting in the light of BCAA uptake regulation being a hallmark of the NCR pathway. Because *SPBPB2B2.01^+^* is strongly induced by NSF, we suggest naming it “NSF-responsive amino acid transporter 1” (*nrt1*^+^). To confirm the specificity of *nrt1*^+^ induction by NSF, we used oleic acid as a negative control for NSF. Oleic acid is a fatty acid with an identical 18 carbon chain and a chemical structure like NSF, but it does not induce adaptive growth (Sun *et al*, 2016). Even with three times the amount of NSF (150 ng/ml), oleic acid did not induce an increase or decrease in the expression of NSF responsive genes (Figure 3B).

**Figure 3.**
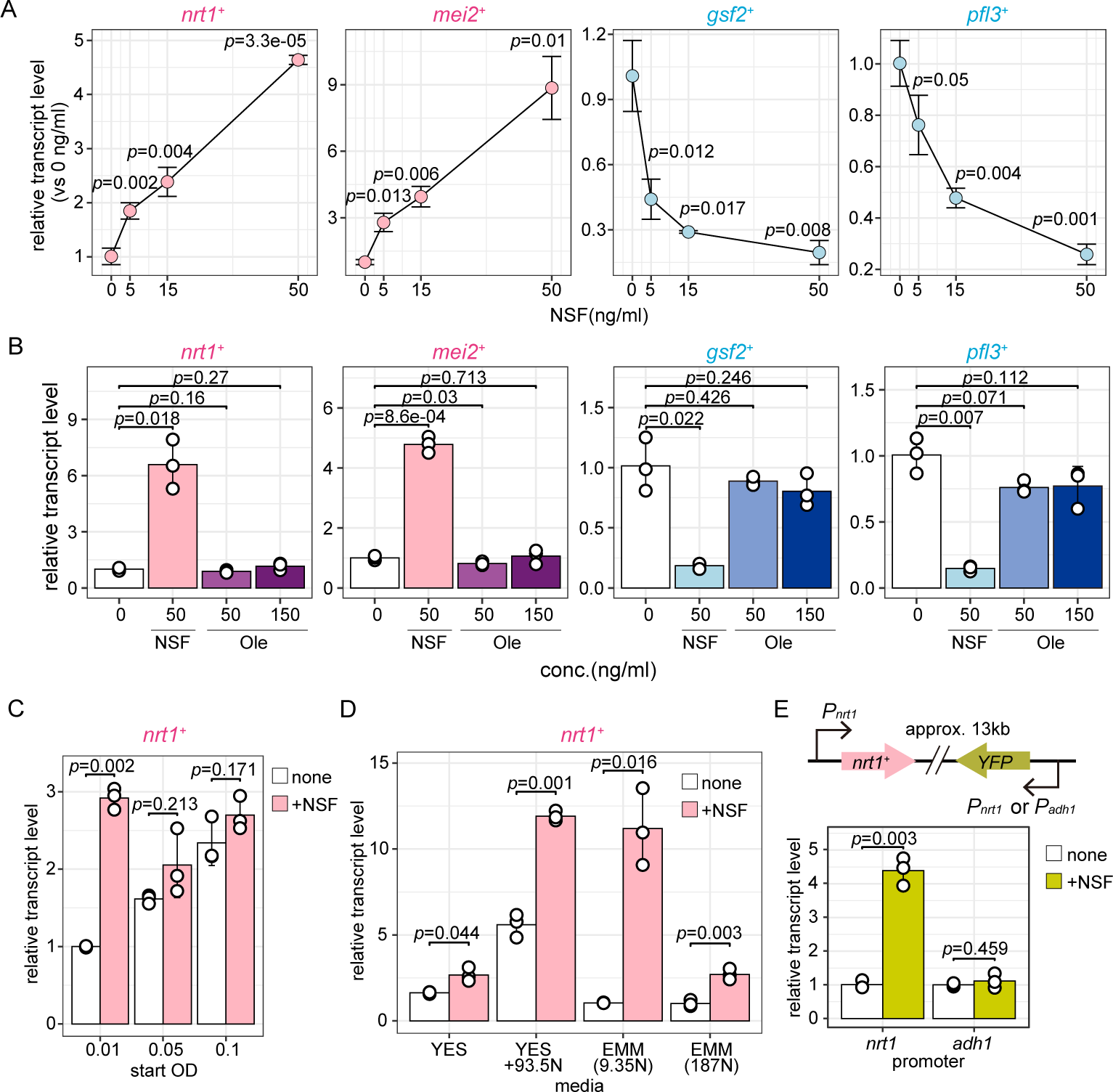
Relative gene expression levels of the two most up- or downregulated genes. **(A)** NSF dose-dependent changes in gene expression. Total mRNA was prepared from cells cultured in EMM(187 mM NH_4_Cl)+Leu+Ade+Ura with increasing concentration of NSF (0, 5, 15, 50 ng/mL) for 6 hours. **(B)** Gene expression changes in response to NSF or oleic acid (Ole) treatment. Total mRNA was prepared from cells cultured in EMM(187mM NH_4_Cl)+Leu+Ade+Ura with NSF (50 ng/mL) or Ole (50, 100 ng/mL) for 6 hours. **(C)** *nrt1^+^*expression in cultures with increasing cell density. Total mRNA was prepared from cells cultured in EMM(187 mM NH_4_Cl)+Leu+Ade+Ura and harvested at different cell densities (OD_600nm_ = 0.01, 0.05, 0.1). **(D)** *nrt1+* expression in YES or EMM medium with various NH_4_Cl concentrations. Total mRNA was prepared from cells cultured in YES, YES+93.5mM NH_4_Cl, EMM(9.35 mM NH_4_Cl)+Leu+Ade+Ura, and EMM(187 mM NH_4_Cl)+Leu+Ade+Ura. **(E)** Yellow fluorescent protein (YFP) gene expression under the control of *nrt1^+^*or *adh1^+^* promoters. The YFP reporter gene was integrated 13 kb from the endogenous *nrt1^+^* locus. Total mRNA was prepared from cells cultured in EMM(187 mM NH_4_Cl)+Leu+Ade+Ura for 6 hours. The mean and standard deviation from three independent experiments are shown. *p*-values were calculated using a two-sided t-test.

We also noted that the stimulatory effect of synthetic NSF on *nrt1*^+^ expression was highest at low cell density (OD 0.01) and ceased at higher concentrations (OD 0.05, 0.1). Yet, expression of *nrt1*^+^ increased with increasing cell density (Figure 3C). This might indicate that *S. pombe* cells produce and secrete NSF, which accumulates with increasing cell density and thereby negating any stimulatory effect of synthetic NSF. Indeed, when grown at low cell density, *nrt1*^+^ expression was induced by NSF irrespective of the nitrogen concentration, or whether cells were grown in rich medium (YES) or synthetic defined minimal medium (EMM) (Figure 3D). Finally, we constructed a strain harboring a YFP gene controlled by either the *nrt1^+^*or *adh1^+^* promoter (Figure 3E). Exposure to synthetic NSF induced YFP expression if driven by the *nrt1^+^*but not the *adh1^+^* promoter. In conclusion, *nrt1^+^*exemplifies an NSF- responsive gene that is controlled at the level of transcription.

### NSF treatment affects transcription factor occupancy on NSF-responsive genes

Coherent with the above finding that NSF triggers changes in gene expression at the level of transcription, our genetic screen revealed eight transcription factor (TF) genes that are required for NSF-mediated adaptive growth. Specifically, these genes encode the TFs Atf1, Adn2, Adn3, Fil1, Hsr1, Php3, Php5, and Reb1 (Table S3). Consistent with a potential role of Atf1 in activating transcription of NSF-responsive genes, we found the GO term “Atf1 activated” enriched among the group of genes that we found to be upregulated in response to NSF treatment (Table S2). *Vice versa*, Adn2 and Adn3 have previously been linked to flocculation and adhesion (Kwon *et al*, 2012), two other GO terms that we found enriched among the genes that are downregulated upon exposure to NSF (Table S2). As mentioned above, *hsr1^+^*was the only gene required for NSF-mediated adaptive growth that was also expressed more upon NSF treatment (Figure 2A). That is, cellular abundance of the TFs Atf1, Adn2, Adn3, Fil1, Php3, Php5, and Reb1 remains unchanged, raising the question how they contribute to changing the transcriptional program. One possibility is that they remain bound to their target genes but become activated or deactivated by NSF directly, or a posttranslational modification, such as phosphorylation in the case of Atf1 (Lawrence *et al*, 2007, 2009a). Alternatively, NSF exposure might strengthen or weaken the binding to their target genes or redirect binding to other genes. To explore this, we tagged Adn2, Atf1, Fil1, Hsr1, Php3, and Reb1 with a 3xFLAG tag at their C-termini and performed chromatin immunoprecipitation coupled to next-generation sequencing (ChIP-seq) with and without NSF treatment. Previously determined target genes of these TFs were significantly enriched in our data set, demonstrating that the experiment has worked (Figure S2A). Globally, TF binding patterns were not grossly altered by NSF treatment. Yet, at a few specific gene promoters we observed a modest increase or decrease in TF enrichment (Figure S2B). These differences in TF occupancy were positively correlated with target gene expression changes. That is, individual genes that were upregulated by NSF tended to be more strongly bound by the TFs, whereas downregulated genes were less occupied by the respective TFs (Figure 4A). For example, increased TF occupancy at the *pex7*^+^ promoter correlated with increased *pex7^+^*mRNA levels, whereas decreased TF occupancy at the *yhb1^+^* promoter correlated with reduced *yhb1^+^* mRNA expression (Figure 4B). This was particularly pronounced for Hsr1 (Figure 4A), which indicates that NSF treatment does not only stimulate *hsr1^+^* expression but also increases as well as reduces Hsr1 binding at specific genes. For Adn2, NSF mainly caused reduced binding at its target genes, which correlated with reduced target gene expression. Occupancy of the other TFs at NSF-responsive genes remained largely the same (Figure 4A and Figure S2B). These results imply that NSF exposure rewires the recipient cell’s transcriptional program, for which the TFs Atf1, Adn2, Adn3, Fil1, Hsr1, Php3, Php5, and Reb1 are indispensable (Table S3).

**Figure 4.**
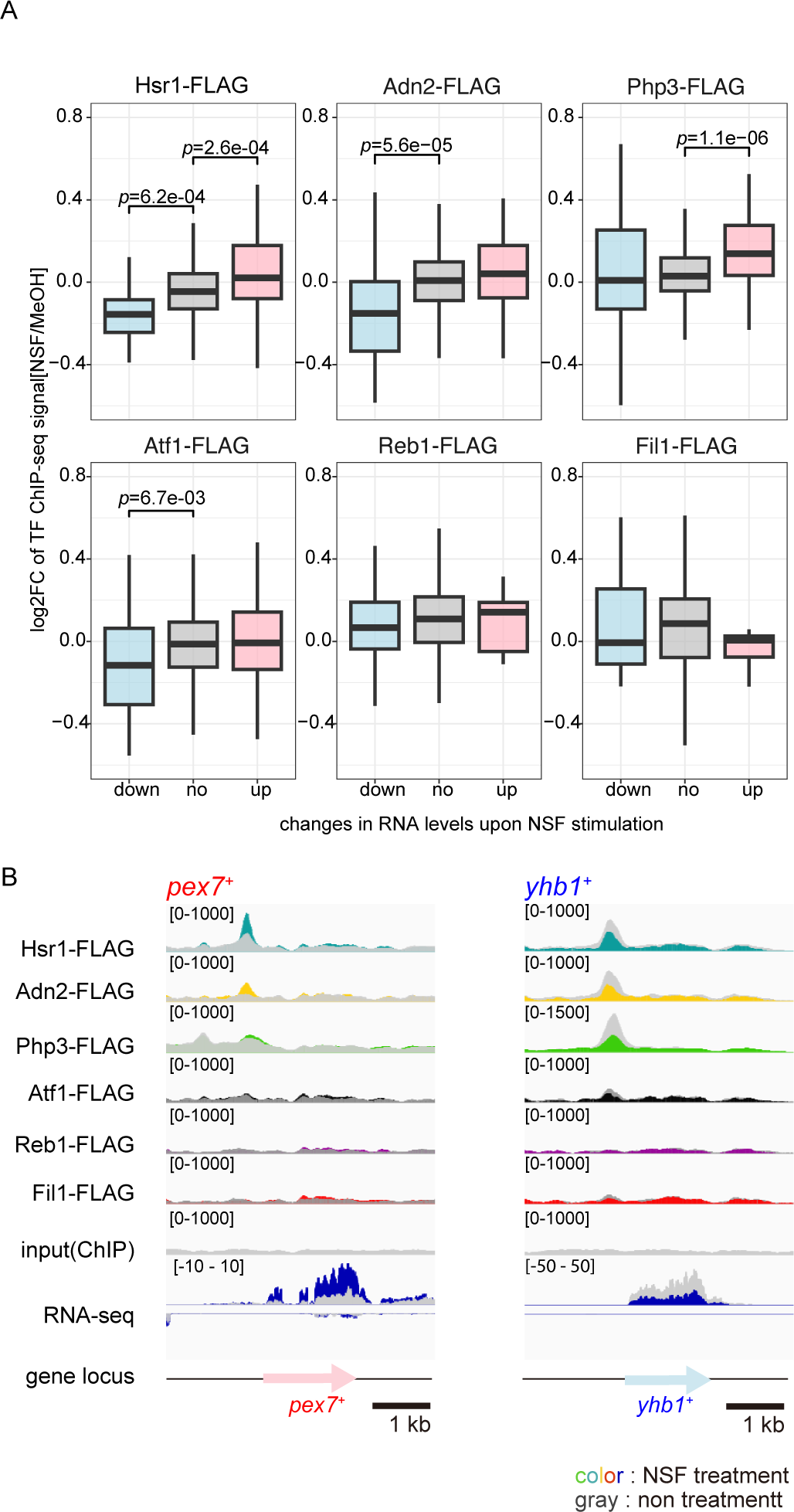
NSF affects occupancy of NSF-linked TFs on NSF-responsive genes. **(A)** Box plots showing log2FC distributions of the respective TF ChIP-seq signals, grouped by NSF-induced gene expression changes shown in Figure 2A (down-regulated, no change, up- regulation). *p*-values were calculated using a two-sided t-test and adjusted using the holm method. **(B)** ChIP enrichments of NSF-linked TFs that were revealed by our screen on the promoters of two representative genes that are either up- or down-regulated by 2 hours NSF treatment (RNA-seq track).

### Ayr1 may metabolize NSF

The foregoing results establish NSF-mediated adaptive growth as a paradigm to investigate how external factors such as chemicals or metabolites in a cell’s environment can lead to transcriptional network changes. To get first insights into the mode of action of NSF, we functionalized NSF with an alkyne tag to generate a probe that can be used for click chemistry, fluorescence microscopy, and affinity purification of putative proteins that might physically interact with NSF (AlkNSF, Figure 5A). Alkyne probes are commonly used in chemical biology because structures and physicochemical properties of small molecules are minimally affected (Wright & Sieber, 2016). Compared to the minimum efficient concentration (MEC) that was previously determined for NSF (12 ng/ml) (Sun *et al*, 2016), we had to use 30-fold higher concentrations of AlkNSF to sustain growth (Figure S3A). Yet, although the MEC of AlkNSF was higher than that of NSF, AlkNSF promoted adaptive growth on EMM(374 mM NH_4_Cl)+Leu plates (Figure 5B) and induced *nrt1*^+^ expression in a dose- dependent manner (Figure 5C). Furthermore, we observed cellular uptake of azide fluor 488- labelled AlkNSF by fluorescence microcopy (Figure S3B). These results indicated that AlkNSF could indeed be successfully employed as a probe in a chemical biology experiment. Therefore, we coupled AlkNSF to azide beads (FG beads®) and incubated these with *S. pombe* whole cell lysates to affinity purify putative NSF interacting proteins. As a control, we preincubated a fraction of the lysate with non-alkylated NSF, which is expected to compete off AlkNSF. Following separation and washing of the AlkNSF-beads, we subjected the immobilized samples to mass spectrometry. This revealed Ayr1 as the sole protein that was co-purifying with AlkNSF and that was competed away with NSF significantly (Figure 5D). Interestingly, *ayr1*^+^ was not among the genes that our screen has identified to be required for adaptive growth (Table S1), which we confirmed in a newly generated *ayr1Δ* strain (Figure 5E). Consistent with this phenotype, NSF-mediated gene expression changes were not abrogated in *ayr1Δ* cells. In contrast, the stimulatory effect of NSF on *nrt1*^+^ or *mei2*^+^ expression was enhanced in the absence of Ayr1 (Figure 5F), suggesting that Ayr1 might function as a negative regulator of NSF-linked changes in gene expression. Because Ayr1 is annotated as a 1-acyldihydroxyacetone phosphate reductase that has been connected to lipid metabolism in yeast (Athenstaedt & Daum, 2000; Ploier *et al*, 2013), it is tempting to speculate that Ayr1 dampens adaptive responses by metabolizing NSF.

**Figure 5.**
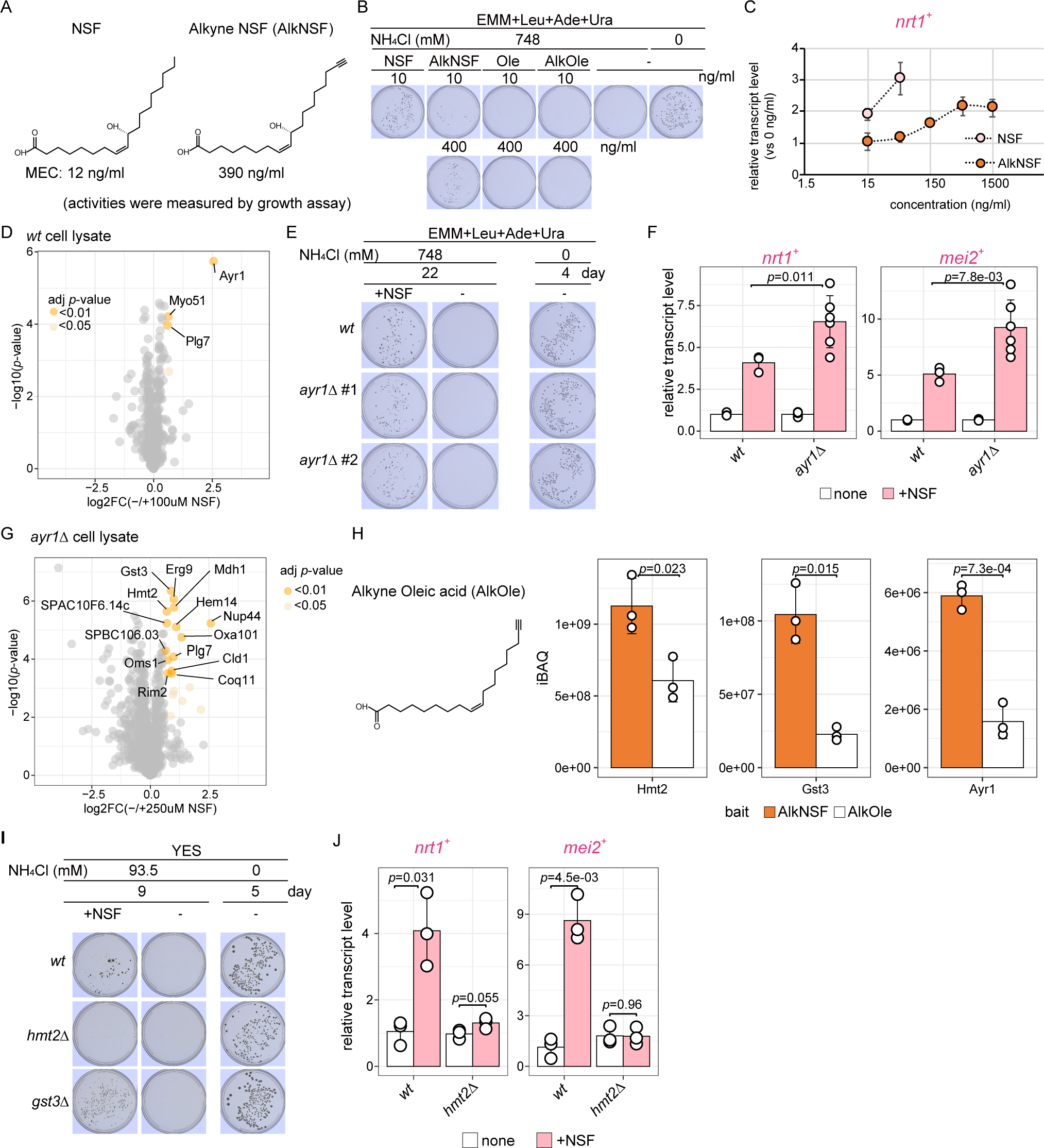
A chemical biology approach identifies potential NSF target proteins. **(A)** Chemical structures of NSF and alkyne-adducted NSF. 10(*R*)-hydroxy-8(*Z*)-octadecenoic acid (NSF) on the left and alkyne-adducted NSF (AlkNSF) on the right. The minimum effective concentration (MEC) was determined by cellular growth assays (see Figure S3A). **(B)** Bioactivity assessments for AlkNSF, Ole, and AlkOle. Leu-auxotrophic cells were spotted onto EMM(374 mM or 0 mM NH_4_Cl)+Leu+Ade+Ura containing either NSF (10ng/mL), alkyne- adducted NSF (AlkNSF) (10 or 400 ng/mL), Ole (10 or 400 ng/mL), or alkyne-adducted oleic acid (AlkOle) (10 or 400 ng/mL). Plates were incubated at 30°C. **(C)** Stimulation of *nrt1^+^*expression by increasing concentrations of NSF (15, 50 ng/mL, lightpink) or AlkNSF (15, 50, 150, 500, 1500 ng/mL, orange) was measured from cells that were grown in EMM(187 mM NH_4_Cl)+Leu+Ade+Ura for 6 hours. Mean and standard deviation from three independent experiments are shown. **(D)** Whole cell lysates were incubated with AlkNSF in the presence or absence of NSF that we used as a competitor. Proteins co-purifying with the AlkNSF probe were identified by mass spectrometry. The data is displayed with enrichment values on the x-axis (AlkNSF versus AlkNSF+NSF) and the *p*-values (moderated t-statistic) on the y-axis. Significantly enriched proteins are labeled by their name. Significance level is indicated by orange (FC >1.5 and adjusted *p*-value <0.01) or light orange color (FC >1.5 and adjusted p- value <0.05). **(E)** NSF-mediated adaptive growth in *ayr1Δ* cells. **(F)** Transcript levels of NSF- responsive genes in *ayr1Δ* cells. Total mRNA was prepared from cells cultured in EMM(187mM NH_4_Cl)+Leu+Ade+Ura. Cells were exposed to 50 ng/mL NSF for 6 hours. **(G)** Whole *ayr1Δ* cell lysates were incubated with AlkNSF. Proteins co-purifying with the AlkNSF probe were identified by mass spectrometry. The data is displayed with enrichment values on the x-axes (AlkNSF versus AlkNSF+NSF) and the BH-adjusted *p*-values (moderated t- statistic) on the y-axis. Significantly enriched proteins are labeled by their name or the systematic gene ID. Significance level is indicated by orange (FC >1.5 and adjusted *p*-value <0.01) or light orange color (FC >1.5 and adjusted p-value <0.05). **(H)** iBAQ values of proteins copurifying with AlkNSF and AlkOle. Whole cell lysates were prepared from wild type cells. *p*-values were calculated using a two-sided t-test. **(I)** NSF-mediated adaptive growth in *hmt2Δ* and *gst3Δ* cells in YES medium. **(J)** Transcript levels of NSF-responsive genes in *hmt2Δ* cells. Total mRNA was prepared from cells cultured in EMM(187mM NH_4_Cl)+Leu+Ade+Ura. Cells were exposed to 50 ng/mL NSF for 6 hours. The mean and standard deviation from three independent experiments are shown. *p*-values were calculated using a two-sided t- test.

### NSF physically and functionally interacts with Hmt2

Because Ayr1 could avert the interaction of AlkNSF with other true interaction partners, we repeated the experiment described above with whole cell lysates generated from an *ayr1Δ* strain. Indeed, this experiment revealed several additional proteins that were significantly enriched compared to the control sample that contained competitive NSF (Figure 5G). Yet, a potential caveat of this result is the hydrophobicity of AlkNSF, which could result in false positive NSF-protein interactions upon prolonged incubation with cell lysates. Therefore, we included an oleic acid alkyne probe (AlkOle), which is similar to AlkNSF but has no biological activity (Figure 5B) and repeated the experiment with *wt* cell lysates to assess which of the above proteins would specifically associate with the AlkNSF but not the AlkOle probe. Reassuringly, Ayr1 was again specifically interacting with AlkNSF. The other proteins that were significantly enriched in the AlkNSF sample were Hmt2 and Gst3 (Figure 5H and Figure S3C). *gst3^+^* was not required for NSF-mediated adaptive growth, neither in our screen nor in growth assays performed with newly generated *gst3Δ* cells (Figure S3D). *hmt2*^+^ was identified by our genetic screen to be required for growth on EMM media (Table S1 and Figure S3D), which can be explained by the cysteine auxotrophy of *hmt2Δ* cells (Pluskal *et al*, 2016). Because our screen was conducted in EMM, *hmt2*^+^ was thus not revealed as a gene necessary for NSF-mediated adaptive growth. Therefore, we reassessed NSF-mediated adaptive growth of *hmt2*Δ and *gst3Δ* cells in YES medium supplemented with 93.5 mM NH_4_Cl. Whereas wild type and *gst3Δ* cells grew in the presence of NSF but not in its absence, *hmt2Δ* cells did not show NSF-mediated adaptive growth (Figure 5I). Consistent with this phenotype, neither *nrt1*^+^ nor *mei2*^+^ expression was induced in the absence of Hmt2 (Figure 5J).

These results identify Hmt2 as a direct target of NSF that is necessary for the activation of NSF-responsive genes. Hmt2 is localized at the inner membrane of mitochondria where it functions as a sulfide:quinone oxidoreductase (SQR) by which hydrogen sulfide acts as an electron donor to the electron transfer chain (ETC) via reduction of a quinone to a hydroquinone (Zhang *et al*, 2021; Weghe & Ow, 1999). Notably, Hmt2 is required for respiratory growth (Malecki *et al*, 2016), which is coherent with our finding that mitochondrial respiration enables evasion of NCR. Thus, the identification of Hmt2 as direct functionally relevant target of NSF provides a first insight into how the NSF signal is received by the recipient cells. How this affects mitochondrial respiration and how this leads to changes in the cell’s transcriptional program are exciting questions that should become subject to future investigations.

## DISCUSSION

Cell-to-cell communication is a widely observed phenomenon among microbes. It is used to react to changes in the environment and to coordinate such responses at the population level to keep the organism thriving. This is often linked to morphological changes such as biofilm formation in bacteria (Miller & Bassler, 2001; Hammer & Bassler, 2003; Mukherjee & Bassler, 2019), mating, or filamentation in fungi (Chen *et al*, 2004; Hornby *et al*, 2001; Merlini *et al*, 2013; Ramage *et al*, 2002). In this study we have focused on cell-to-cell communication mediated by NSF, a diffusible oxylipin that is produced and secreted by *S. pombe* cells to their extracellular milieu. In contrast to other communication systems, NSF does not induce noticeable changes in *S. pombe*’s morphology. A characteristic effect of NSF is that it enables the uptake of BCAA, which is essential for cells that cannot produce their own BCAA. Because excess ammonium supplementation in the media leads to the inhibition of BCAA uptake, a phenomenon known as nitrogen catabolite repression or NCR (Mitsuzawa, 2006; Magasanik & Kaiser, 2002; Ljungdahl & Daignan-Fornier, 2012), it is generally assumed that NSF functions to control the usage of different nitrogen sources (Takahashi *et al*, 2012). However, the pathways and mechanisms by which NSF functions and whether these are restricted to nitrogen metabolism have remained elusive. We have performed a genetic screen with a non-essential gene deletion library to identify factors that are required for NSF-mediated adaptive growth at high ammonium concentrations. Essentially, the screen was designed such that every mutation that disables the uptake of a BCAA would score. This has revealed 117 mutants that grew normally without ammonium, but not at high ammonium concentrations despite the addition of synthetic NSF and the presence of exogenous BCAA. Interestingly, many of the genes identified in this screen are required for mitochondrial respiration (Figure 1C), suggesting that NSF-mediated adaptation to high ammonium concentrations depends on the respiratory capacity of the cell. This hypothesis is further supported by our observation that inhibition of mitochondrial electron transport by antimycin A stops cells from growing in the presence of high ammonium concentrations. Conversely, indirect induction of respiration by glucose limitation makes NSF dispensable for growth. Moreover, we found that NSF triggers a respiration-like gene expression pattern, and that NSF physically interacts with the mitochondrial sulfide:quinone oxidoreductase Hmt2. Therefore, we propose a model in which NSF activates mitochondrial respiration, which eventually leads to global changes in gene expression, the evasion of NCR, and the uptake of BCAA. The latter is reminiscent of previous work conducted in budding yeast, which revealed that the induction of respiration by glucose de-repression demands and increases the uptake of leucine for intermediates of the TCA cycle (Hothersall & Ahmed, 2013).

We find it interesting that NSF triggers a change from a fermentation- to a respiration-like gene expression program independently of the carbon source. This raises the question how *S. pombe* could benefit from respiration in the presence of glucose, which is well known to promote rapid proliferation by fermentation (Pfeiffer & Morley, 2014; Hagman & Piškur, 2015). Importantly, the concentration of NSF in the milieu is positively correlated with cell density. That is, the fewer cells the less NSF will be secreted. Therefore, the switch from fermentation to respiration would be expected to occur only when cells become denser and the concentration of NSF increases. Indeed, *nrt1*^+^ expression increased with higher cell density without the addition of extra NSF (Figure 3B). Moreover, the stimulatory effect of synthetic NSF was highest at low cell density and ceased when the population grew denser. This raises the possibility that NSF may function as a rheostat to prepare a population of *S. pombe* cells for a foreseeable shortage of glucose when they grow exponentially. Thereby, reaching a certain NSF concentration would cause a switch to respiration, also known as diauxic shift (Brauer *et al*, 2005; Dickinson & Schweizer, 2019; Bartolomeo *et al*, 2020), preparing the cells to utilize other carbon sources such as ethanol, because glucose will eventually become limiting. Whereas diauxic shifts are well-established, to our knowledge this phenomenon has not been linked to cell-to-cell communication. Because NSF is an intraspecies-specific signal (Yashiroda & Yoshida, 2019; Takahashi *et al*, 2012; Sun *et al*, 2016), the NSF-mediated shift from fermentation to respiration would exemplify a “social interaction” between cells of the same species, preparing them for an imminent shortage of their preferred carbon source. This would increase the population’s fitness and thus confer *S. pombe* a competitive advantage in such environment. In that sense, NSF would not strictly revoke NCR only, but also CCR, and it could thus be considered as a general mediator of catabolite repression.

An important question that will need to be addressed in the future is how NSF exposure triggers the observed change in *S. pombe*’s gene expression program. Because our chemical biology approach has not revealed transcription factors or chromatin regulators as direct targets of NSF, it is unlikely signaling directly to the nucleus. Rather, identification of Hmt2 as a physical interaction partner of NSF suggests that the transcriptional changes observed in the nucleus are a consequence of altered metabolic activity of mitochondria. Hmt2 is a sulfide:quinone oxidoreductase (SQR), which couples sulfide oxidation to coenzyme Q_10_ reduction in the ETC. This reaction generates highly reactive sulfur species (RSS) such as glutathione persulfide, which can be further oxidized or can modify cysteine residues in proteins by persulfidation (Filipovic *et al*, 2018; Cuevasanta *et al*, 2017; Mishanina *et al*, 2015; Mustafa *et al*, 2009a, 2009b). Interestingly, there are several reports that persulfidation of transcription factors could regulate their activities (Yao *et al*, 2023; Tian *et al*, 2021; Shimizu *et al*, 2023). Thus, persulfide signaling is an attractive hypothesis for how NSF signaling could be relayed to the cell nucleus that will be worthwhile testing.

## MATERIALS AND METHODS

### Yeast strains and growth media

*S. pombe* strains were generated following a PCR-based protocol (Bähler *et al*, 1998) and strains were validation by colony PCR. For a list of strains generated in this study see Table S4. Cells were grown in rich yeast extract (YE) medium with 3% glucose, YE with 2 mM each adenine and uracil (YES), or in minimal (EMM) medium with 2% glucose and 0.5% (93.5 mM) ammonium chloride (NH_4_Cl). Appropriate leucine, adenine, uracil and glutamate (2 mM) were added to EMM medium. For nitrogen catabolite repression (NCR) condition, EMM was supplemented with extra NH_4_Cl (final concentration 187∼748mM) and for non NCR condition, EMM without ammonium sulfate (0 mM NH_4_Cl) with 2 mM Leu as a sole nitrogen source. For supplementation of NSF, 100 µL of 10 µg/mL synthesized NSF (10(*R*)-hydroxy- 8(*Z*)-octadecenoic acid) (Sun *et al*, 2016) dissolved in 50% methanol was added onto 20 mL agar media (final concentration 50 ng/mL). Where indicated, media were supplemented with antimycin A (0.4 µM) for inhibition of respiration or with lower glucose (0.08%) for induction of respiration. For cell mating, sporulation agar (SPA) medium was used.

### Plasmids

Plasmids were cloned by standard molecular biology techniques. For a list of plasmids generated in this study see Table S5.

### Genetic screening for adaptive growth

To enable selection for the *h^-^* mating type in the Leu-auxotrophic knock-out library, an *h^-^* strain having the drug-resistant markers at the *mat1* locus was generated. The kanamycin resistance (kanMX) marker was inserted downstream of the *mat1-Mc* gene, which was subsequently replaced with the nourseothricin resistance marker (natMX) by the marker switch technique. First, two homology arms were amplified from genomic DNA by PCR using two primer sets: insert-kanR-F1 and insert-kanR-F2, and insert-kanR-R1 and insert-kanR-R2. The purified PCR products were then used for assembling the disruption fragment by amplifying the kanMX cassette by PCR. The resultant PCR products were introduced into the JY265 strain. The strain in which the kanMX marker was replaced by the natMX marker were generated as previously described (Sato *et al*, 2005). We prepared the PCR product using MS-TEP and MS-TET as primers and pCR2.1-nat as a template. The Leu-auxotrophic knock- out library (3225 mutants) was made from the BIONEER haploid deletion mutant library v5.0 (3420 mutants) by mating with *leu1-32* auxotroph on SPA+Ade+Ura+Leu (2 mM each) medium. For selection after mating, mutants were grown on SD+Leu media, twice on YE + G418 and nourseothricin, twice. For the first screening, mutants are spotted onto 48-well plate solid EMM (748 mM NH_4_Cl) +Leu (2 mM) media with or without a prototroph spotted next to mutant spot. Plates were incubated at 30 °C, and images were acquired after 5 or 6 days. For the second screening, mutants were spotted onto 48-well plate solid EMM (0 mM or 748 mM NH_4_Cl) +Leu media with NSF or methanol. Plates were incubated at 30 °C, and images were acquired after 5 or 6 days. GO category enrichments in each mutant cluster were calculated using the AnGeLi web tool (Bitton *et al*, 2015)

### Oxygen consumption rate measurement

The respiratory capacity was assessed using the Seahorse XF HS mini analyzer (Agilent Technologies). Seahorse XFp cell culture miniplate was coated with 50 μg/mL poly-lysine (50 µL each well) for 30 min at room temperature and then aspirated followed by air drying at 4 °C overnight. The Seahorse XFp extracellular flux cartridge was hydrated with sterile water and incubated overnight at 30 °C and Seahorse XF calibrant solution (Agilent Technologies). After this, the water was removed, then added the calibrant solution to the sensor cartridge and incubated at 30 °C for 1 hour before measurement.

Yeast cultures were grown in YES media at 30 °C for 16 hours, followed by subculturing in EMM(561 mM NH_4_Cl)+Leu+Ade+ura media with or without NSF (50 ng/mL) for 30 °C for 8 hours, start OD_600nm_ at 0.01. For measurement, all cultures were diluted in EMM(561 mM NH_4_Cl)+Leu+Ade+ura to seed an OD_600nm_ at 0.01 into a poly-lysine coated Seahorse XFp cell culture miniplate in 50 μL of Seahorse XF assay media (Agilent Technologies). For background measurement, two wells containing only assay media were included. The loaded plate was centrifuged at 300 x *g* for 3 min at room temperature. After centrifugation, the volume of the media was made up to 150 μL, and the loaded plate was incubated for 30 min at 30 °C to facilitate the transition of the plate into the Seahorse machine’s temperature. Measurements by Seahorse were performed for 10 cycles.

### RNA isolation and cDNA synthesis

Total RNA was isolated using the MasterPure Yeast RNA Purification Kit (Epicentre). cDNA was synthesized using the PrimeScript RT Master Mix (Takara).

### Total RNA sequencing and analysis

Total RNA libraries were prepared with TruSeq Stranded Total RNA kit (Illumina) according to the manufacturer’s instructions and sequenced with an Illumina HiSeq2500 (50 bp single-end). RNA-seq reads were aligned to the S. pombe genome (ASM294 version 2.24) using STAR (Dobin *et al*, 2013) (version 2.7.3a) (STAR—runMode alignReads—outFilterType BySJout—outFilterMultimapNmax 100—outFilterMismatchNoverLmax 0.05—outSAMmultNmax 1—outMultimapperOrder Random—outSAMtype BAM SortedByCoordinate—outSAMattributes NH HI NM MD AS nM—outSAMunmapped Within). The reads per gene were counted with featureCounts (Liao *et al*, 2014) of uniquely mapping reads only (useMetaFeatures = TRUE, allowMultiOverlap = FALSE, minOverlap = 5, countMultiMappingReads = FALSE, fraction = FALSE, minMQS = 255, strandSpecific = 2, nthreads = 20, verbose = FALSE, isPairedEnd = FALSE). An external feature annotation file for *S. pombe* was used based on a GFF3 file from PomBase (Lock *et al*, 2018) that was converted to a GTF file with rtracklayer (Lawrence *et al*, 2009b). Differential gene expression analysis was performed using DESeq2 (Love *et al*, 2014). Each MA plot depicts log2 fold changes of cells with NSF treatment against cells without NSF treatment. For GO term enrichment analysis, we downloaded GO term annotation for *S. pombe* genes from Pombase and used the GO.db R package to build a GO annotation map for *S. pombe*. Then we used the enricher function from the ClusterProfiler package to test for enriched GO terms in up or down regulated genes compared to all tested genes. Total RNA sequencing data have been deposited at the NCBI Gene Expression Omnibus (GEO) database and are accessible through GEO series number GSE250095.

### RT–PCR

PCR on cDNA was performed using the fast-cycling PCR kit (Qiagen). Primer sequences are listed in Table S6.

### Chromatin immunoprecipitation (ChIP) and ChIP-sequencing

ChIP experiments were performed as described previously (Kuzdere *et al*, 2023) with 2.5 µg anti-FLAG M2 antibodies (Sigma). ChIP-sequencing libraries were generated using the NEBNext Ultra II DNA Library Prep Kit (New England Biolabs) and sequenced with an Illumina HiSeq2500 (50 bp single-end). Raw data were demultiplexed, converted to fastq format using bcl2fastq2 (v1.17), and mapped using STAR (Genome_build: Spombe.ASM294v2.24). For bigwig track generation by bedtools (v2.26.0) and bedGraphToBigWig (from UCSC binary utilities), nonaligned reads were discarded and read coverage was normalized to 1 million genome mapping reads (RPM). Peak finding was done using MACS2 (Gaspar, 2018) and peaks with a score above 100 were used. Differential binding was calculated using Limma – Voom and the total number of mapped reads as library size. To find the corresponding gene promoters under TF binding peaks, we used R package GenomicFeatures and the “nearest” function of GenomicRanges. *S. pombe* genomic range file was downloaded from Pombase (Lock *et al*, 2018) and non-coding RNA genes were removed from the list. And then, genomic ranges of protein coding gene were changed to promoter range (200 bp upstream from transcription start site).

### Microscopic observation of AlkNSF with click chemistry

Cells were inoculated into EMM (187mM NH_4_Cl)+Leu+Ade+Ura, with 5 µM AlkNSF (manuscript in preparation) adjusted to OD_600nm_ at 0.01. The culture was incubated at 30 °C for 4 hours. Subsequently, cells were harvested and fixed in 70% ethanol. After fixation, cells were washed twice with TBS buffer, followed by incubation in a solution containing 20 µM Azide fluor (Sigma), 2 mM CuSO_4_ (Sigma), and 10 mM ascorbate in TBS buffer in the dark for 30 min. Following three washes with TBS, cells were observed under a fluorescence microscope.

### Affinity purification of AlkNSF

To make AlkNSF pre-coupled beads, 2.5 mg (125 µL) azide beads (Tamagawa Seiki) with 125 µM AlkNSF or Alkyne oleic acid (AlkOle) (Cayman), 62.5 µM tris[(1-benzyl-1H-1,2,3-triazol-4- yl)methyl]amine (TBTA) (Sigma), 1.25 mM CuSO_4,_ and 1.25 mM (+)-sodium L-ascorbate (Sigma) were incubated for 16 hours at room temperature. The pre-coupled beads were three-time washed with t-BuOH(Supelco)/DMSO/water (4:1:5) and three-time washed with 50% (v/v) methanol. The washed beads were washed and resuspended with Lysis buffer (150 mM NaCl, 20 mM HEPES pH 7.5, 5 mM MgCl_2_, 1 mM EDTA pH 8.0, 10% glycerol, 0.25% (v/v) Triton X-100, 0.5 mM DTT, and 1x HALT protease inhibitor cocktail (Thermo Fisher Scientific)).

Cells grew in YES media for 16 hours and were harvested at 2500 rpm. Harvested cells were washed with TBS (50 mM Tris-HCl pH 7.5, 150 mM NaCl) twice. Cells were resuspended in Lysis buffer and were disrupted with silica beads by bead-beating machine (MP Biomedicals™ FastPrep-24™ 5G Instrument, 3x 20s at 6.5m/s, 3min breaks on ice in between rounds). Cell lysates were collected from tube having punched a hole in the bottom with a needle followed by centrifugation. Crude lysates were centrifugated with 13000 rpm for 10 min at 4 °C. Protein concentration of clear lysates was measured by Bradford assay. As competition assay, cell lysates were incubated with NSF (100 µM for parent cell lysate or 250 µM for *ayr1Δ* cell lysate) for 2 hours at 4 °C. Pre-coupled beads of AlkNSF or AlkOle were added and were incubated for 2 hours at room temperature. The beads were washed twice each with lysis buffer and wash buffer (100 mM NaCl, 20 mM HEPES pH 7.5, 5 mM MgCl_2_, 1 mM EDTA pH 8.0, 10% glycerol, 0.25% (v/v) Triton X-100). Washed beads were resuspended with 6 µL digest buffer (3 M guanidine HCl, 20 mM HEPES pH 8.5, 10 mM CAA, 5 mM TCEP) and supplemented with 0.2 µg LysC. After incubation for 4 hours at room temperature, 17 µL of 50 mM HEPES pH 8.5 and 20 ng trypsin were added to further digested the proteins. Trypsin digestion was conducted overnight at 37 °C.

### Mass spectrometry

The generated peptides were acidified with TFA to a final concentration of 0.8% and analyzed by LC–MS/MS on an Orbitrap FUSION LUMOS or ECLIPSE tribrid mass spectrometer (Thermo Fisher Scientific) connected to a Vanquish Neo UHPLC (Thermo Fisher Scientific) with a two-column set-up. The peptides were applied onto a C18 trapping column in 0.1% formic acid and 2% acetonitrile in H_2_O. Using a flow rate of 200 nL/min, peptides were separated at RT with a linear gradient of 2–6% buffer B in buffer A in 1 min followed by a linear increase from 6 to 20% in 42 min, 20–35% in 22 min, 35–45% in 2 min, 40–100% in 1 min, and the column was finally washed for 10 min at 100% buffer B in buffer A (buffer A: 0.1% formic acid; buffer B: 0.1% formic acid in 80% acetonitrile) on a 15 cm EASY-Spray column (Thermo Fisher Scientific) mounted on an EASY-Spray^TM^ source (Thermo Fisher Scientific). The survey scan was performed using a 120,000 resolution in the Orbitrap followed by an HCD fragmentation of the most abundant precursors. The fragments mass spectra were recorded in the ion trap according to the recommendation of the manufacturer (Thermo Fisher Scientific). Protein identification and relative quantification of the proteins was performed with MaxQuant v.2.2.0.0 using Andromeda as search engine (Cox *et al*, 2011), and label-free quantification (LFQ) (Cox *et al*, 2014). The fission yeast subset of the UniProt v.2021_05 combined with the contaminant database from MaxQuant was searched and the protein and peptide FDR were set to 0.01. The LFQ intensities estimated by MaxQuant were analyzed with the einprot R package (https://github.com/fmicompbio/einprot) v0.7.6 as previously described (Welte *et al*, 2023).

### Data availability

All custom codes used to analyze data and generate figures are available upon reasonable request.

RNA-Seq and ChIP-seq data sets have been deposited to the Gene Expression Omnibus with the data set identifier GSE250095. The mass spectrometry proteomics data have been deposited to the ProteomeXchange Consortium via the PRIDE (Perez-Riverol *et al*, 2021) partner repository with the dataset identifier PXD047795.

## ACKNOWLEDGEMENTS

We thank the members of the Bühler lab for their constant support and discussions. Special thanks go to Fabio Mohn for data deposition, and Yukiko Shimada and Nathalie Laschet for technical support. We are also grateful to the FMI Functional Genomics facility for library construction and next-generation sequencing and to Laurent Gelman for assistance with fluorescence microscopy. This work was supported by the Novartis Research Foundation (M.B.), the Japan Society for the Promotion of Science (JSPS; KAKENHI Grant Numbers 18H02131 and 22K05397 to Y.Y. and G.H., 19K15755 to S.O, and 23H05473 and 23H04882 to M.Y.) and the Ohsumi Frontier Science Foundation (to Y.Y.).

## AUTOR CONTRIBUTIONS

**Marc Bühler:** Conceptualization; resources; supervision; funding acquisition; visualization; project administration; writing—original draft, review and editing. **Yoko Yashiroda**: Conceptualization; resources; supervision; funding acquisition; project administration; writing—review and editing. **Shin Ohsawa**: Conceptualization; formal analysis; validation; investigation; visualization; methodology; funding acquisition; writing—original draft; writing—review and editing. **Vytautas Iesmantavicius:** Formal analysis; investigation. **Michaela Schwaiger**: Formal analysis; investigation, **Rio Hashimoto**: Formal analysis; investigation. **Hiromitsu Moriyama**: Conceptualization; supervision; writing—review and editing. **Hiroaki Matoba**: Conceptualization; methodology; writing—review and editing, **Go Hirai**: Conceptualization; methodology; funding acquisition; writing—review and editing. **Mikiko Sodeoka**: Conceptualization; supervision; writing— review and editing. **Atsushi Hashimoto**: Formal analysis; investigation. **Akihisa Matsuyama**: Formal analysis; investigation. **Minoru Yoshida**: Conceptualization; resources; supervision; funding acquisition; project administration; writing—review and editing

---

**Figure S1. Differential gene expression analysis of cells treated with NSF for 4 and 6 hours.** MA plot showing differential gene expression (log2FC) in cells treated with MeOH or NSF for 4 and 6 hours (y-axis). The 2-hour treatment is shown in Figure 2A. x-axis denotes total transcript abundance in counts per million (cpm) in both conditions. *p*-values were calculated using the wald test and adjusted with the Benjamini and Hochberg method. Up- or down-regulated genes (FC >1.5 and adjusted *p*-value <0.05) are highlighted in pink or blue, respectively. NSF-linked genes revealed by the genetic screen are marked with a black outline. The names of the top two upregulated and downregulated genes are written red and blue, respectively. Green color was used to label the genes encoding transcription factors.

**Figure S2. ChIP-seq analysis of TFs required for NSF-mediated adaptive growth. (A)** ChIP enrichments of Hsr1-FLAG, Adn2-FLAG, Php3-FLAG, Atf1-FLAG, Reb1-FLAG, and Fil1-FLAG on NSF-linked genes. RNA-seq tracks show expression changes after 2 hours NSF treatment (gray versus blue track). **(B)** Changes in ChIP-seq signal of the respective transcription factors are shown in MA plots. Log2 fold-changes are shown on the y-axis. *p*- values were calculated using the wald test and adjusted with the BH method. The x-axis shows the average normalized read counts (log10) of samples with and without NSF treatment. Significantly changing ChIP-seq signals (FC >|1.5| and adjusted *p-*value <0.05) are highlighted in orange.

**Figure S3. Finding NSF interacting proteins with a chemical biology probe. (A)** Determination of the minimum effective concentration of AlkNSF. *eca39Δ* cells were spotted onto EMM(Glu)+ILV+Ade+Ura with decreasing AlkNSF concentrations. As a positive control for this assay (PC), cells were exposed to secreted signaling factors that were purified from the supernatant of an *S. pombe* culture with the ethyl acetate method (Sun *et al*, 2016). 50% methanol (MeOH) served as a negative control. The plate was incubated for 6 days at 30 °C. **(B)** Visualization of cellular AlkNSF uptake by fluorescence microscopy. Cells were incubated with 5 µM AlkNSF in EMM(187 mM NH_4_Cl)+Leu+Ade+Ura for 4 hours. Subsequently, AlkNSF was conjugated with azide-flour 488 by click chemistry. **(C)** iBAQ values of proteins that co-purify with AlkNSF and AlkOle probes when incubated with wild type cell lysates. *p*-values were calculated using a two-sided t-test. **(D)** NSF-mediated adaptive growth of *hmt2Δ* or *gst3Δ* cells.

